# Compressive strengths of PEG gels with glycerol and bioglass particles

**DOI:** 10.1101/405779

**Authors:** Ariel Golshan, Jenesis A. Curtis, Vasilios Lianos, Sina Y. Rabbany, Roche C. de Guzman

## Abstract

Poly(ethylene glycol) (PEG)-based materials can potentially be used as biomechanical matrices for regenerative medicine implants including the replacement of intervertebral (IV) discs. Glycerol and other plasticizers (low-MW PEG, propylene glycol, and sorbitol) were added to the bulk PEG matrix, gelled using chemical and photochemical methods at different temperature and pressure settings, and compression properties acquired and analyzed. Incorporation of surface bioactive glass particles shortened the blood clotting time, while alginate and laponite additives improved the gel’s mechanical properties to 645 kPa compressive modulus, 12% yield strain, and 79 kPa yield strength. This IV disc-modeled system endured the cyclic loading and unloading test indicative of an elastic response; but required improvement of its biomechanical tolerance.

## I. INTRODUCTION

Poly(ethylene glycol) (PEG) is a commonly-used biomaterial due to its proven safety record, nontoxicity, and biocompatibility.^1-5^ However, as tissue engineering materials specifically for intervertebral (IV) disc applications, PEG hydrogels alone are not sufficient due to their inherent fragility.^6-9^ Plasticizers are low-molecular weight (MW) compounds that increase the flexibility of polymers by filling-up the network spaces, allowing increased polymer mobility.^10-12^ In this study, to improve the compressive strengths and other important mechanical properties of PEG hydrogels, U. S. Food and Drug Administration (FDA)’s generally recognized as safe (GRAS) substances: glycerol (glycerin), propylene glycol, and sorbitol, and relatively small-MW uncrosslinked PEG were utilized as plasticizers (Table I). The crosslinking process of the bulk PEG matrix was conducted using PEG diacrylate (PEGDA) chain intermediates, and the effect of chemical versus photochemical PEGDA chain elongation leading to gelation, was also investigated at various temperature and pressure conditions, as they (type of crosslinkers, temperature, and pressure) affect the physical properties of gels.^13-17^

**Table I.**
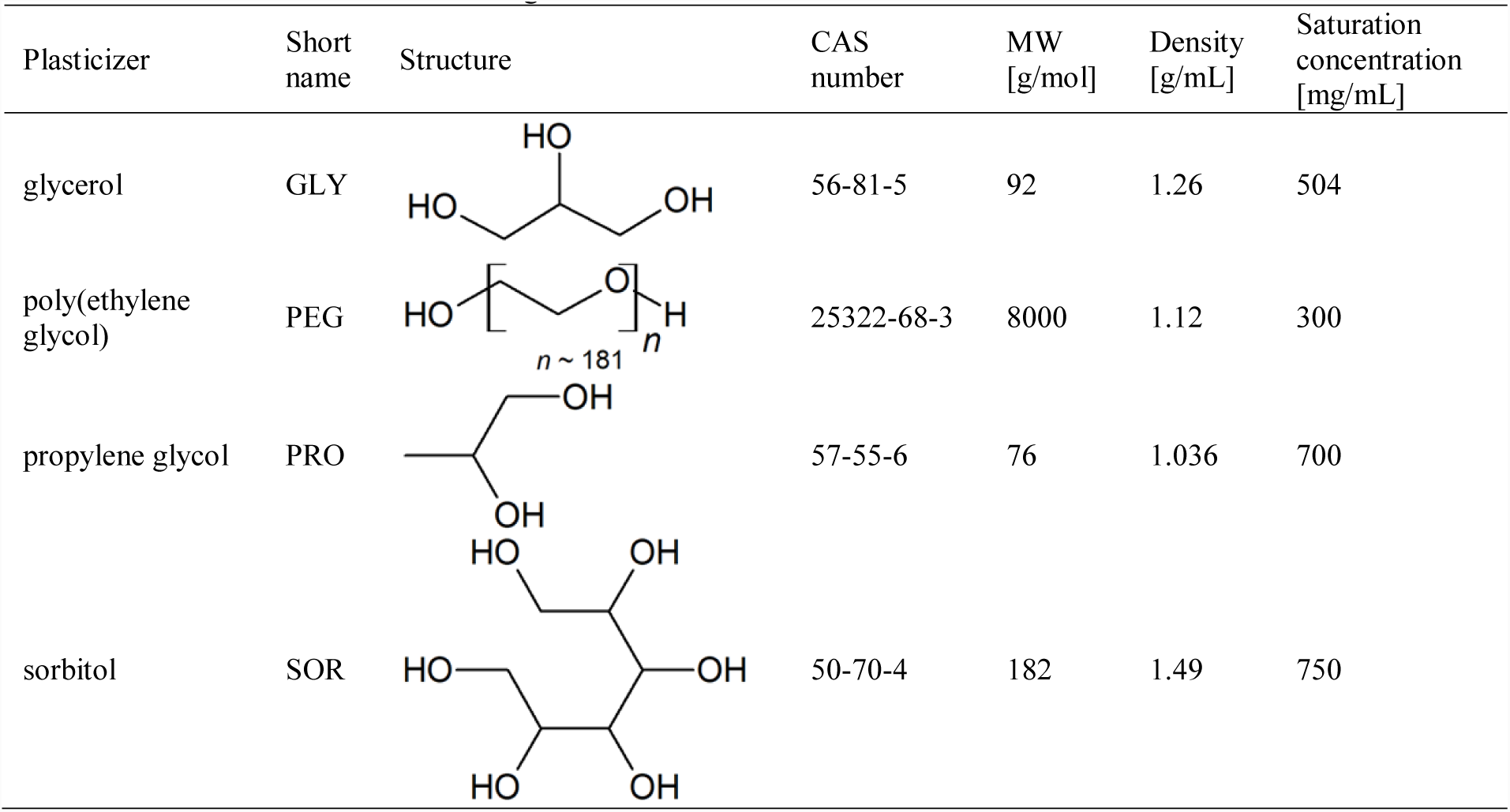
Plasticizers used in bulk PEG gels.

| Plasticizer | Short name | Structure | CAS number | MW [g/mol] | Density [g/mL] | Saturation concentration [mg/mL] |
| --- | --- | --- | --- | --- | --- | --- |
| glycerol | GLY |  | 56-81-5 | 92 | 1.26 | 504 |
| poly(ethylene glycol) | PEG |  | 25322-68-3 | 8000 | 1.12 | 300 |
| propylene glycol | PRO |  | 57-55-6 | 76 | 1.036 | 700 |
| sorbitol | SOR |  | 50-70-4 | 182 | 1.49 | 750 |

Alginate biomaterials, the naturally-derived polysaccharides that gel via chain interactions linked by multivalent cations (such as Ca^2+^), have been utilized for IV-disc regeneration research.^8, 18, 19^ Laponite is a mixture of inorganic minerals with two-dimensional (2D) disk-shaped architecture for the improvement of bulk mechanical properties.^20^ Together, these additives (alginate and laponite) were incorporated in the PEG gel assembly to potentially sustain the loads experienced by native lumbar IV discs.^8, 18, 21^ Finally, bioactive glass or bioglass^22^ particles, the engineered slowly-degrading materials known to bind to bones without inducing a foreign-body capsule reaction, were synthesized in the lab then layered onto the surfaces of the cylindrical gel and evaluated for blood clotting to imply host tissue integration.

## II. EXPERIMENTAL

### A. Materials

The following chemicals were purchased from Sigma-Aldrich (St. Louis, MO): poly(ethylene glycol) diacrylate (PEGDA; MW = 700 g/mol), glycerol (GLY; MW = 92 g/mol), poly(ethylene glycol) (PEG; MW = 8000 g/mol; n ∼ 181), propylene glycol (PRO; MW = 76 g/mol), sorbitol (SOR; MW = 182 g/mol), ammonium persulfate (APS), tetraethylethylenediamine (TEMED), methanol (MeOH), phosphate-buffered saline (PBS), silica (SiO_2_), sodium oxide (Na_2_O), calcium oxide (CaO), phosphorus pentoxide (P_2_O_5_), calcium chloride (CaCl_2_), and sodium alginate (ALG; product # W201502). Irgacure^®^ 2959 (I2959) was obtained from BASF (Vandalia, IL) and Laponite^®^ RD (LAP; synthetic layered silicate made from SiO_2_, MgO, Li_2_O, and Na_2_O) from BYK Additives (Gonzales, TX). Deionized water was used as the solvent for preparing solutions unless stated otherwise.

### B. Gel synthesis using photochemical-crosslinking with varying plasticizers

Cylindrical gels were prepared into wells of 24-well poly(styrene) plates at final concentrations of: 20% (V/V) PEGDA, 10 mg/mL I2959 (diluted from 100 mg/mL I2959 in MeOH), and varying amounts of plasticizers (GLY, PEG, SOR, and PRO) up to their saturation limits. The saturation limits of respective plasticizers in gels were determined by adding increasing amounts of plasticizers (10% to 75%) until the gel could no longer hold the plasticizer (phase separation observed). The maximum amounts of gel-incorporated plasticizers were found to be: GLY at 40% (V/V) = 0.4 mL/mL = 504 mg/mL, PEG at 30% (m/V) = 300 mg/mL, PRO 70% (m/V) = 700 mg/mL, and SOR at 75% (m/V) = 750 mg/mL (Table I). These values were then set to be the 100% relative saturation levels. PEG, SOR, and PRO were prepared at higher concentrations prior to dilutions. After mixing the components into uniform solutions, samples were photo-crosslinked by exposure to a 36-W, 254-nm UV light for 20 minutes at ambient temperature (21 °C).^23^

### C. Gel compression analysis

Strength and viscoelastic properties of gels were characterized through unconfined compression until fracture. Freshly-made gels were biopsy-punched to obtain cylindrical specimens with diameter (d) = 11 mm, loaded onto a platen, then compressed with a parallel blunt probe at a speed of 0.1 mm/s using the Instron 3345 mechanical tester (Norwood, MA) with a 100-N load cell. Settings were configured and data (relative displacement versus force) were acquired with the Bluehill 3 software (Instron). Data were processed in Excel (Microsoft, Redmond, WA) and MATLAB (Mathworks, Natick, MA) to obtain true stress and strain results.^24^ Compression properties: elastic modulus at compression (E_c_), and yield and ultimate strains and stresses were determined.

### D. Gel synthesis using chemical crosslinking at different temperatures and pressures

PEG-based gels were fabricated via chemical crosslinking by homogenously mixing the following: 20% (V/V) PEGDA, 30% (V/V) GLY plasticizer, 1% (V/V) TEMED, and 1 mg/mL APS into the chamber of a heated die set (Across International, Livingston, NJ). A 10 mg/mL APS intermediate solution was prepared and used prior to the gelation reaction. Gels (d = 25 mm and h = 5 mm) were allowed to set at different temperatures: 21, 37, 59, and 80 °C using a temperature controller for 15 minutes, equilibrated back to room temperature (RT), biopsy-punched at d = 8 mm, and unconfined compression testing conducted as described above. Gels without the GLY plasticizer were utilized as controls. Representative PEG-GLY gels were also fabricated under 90 MPa of gauge pressure (compared to gels made in atmospheric pressure) for 15 minutes at 21°C within an MP15 desktop pellet press (Across International).

### E. Glycerol release assay

The stability of GLY in PEG-GLY gels were tested by spectrophotometry. Hydrogels (20% PEGDA and 30% GLY) were made similar to the abovementioned protocol using chemical crosslinkers. 8-mm diameter samples were obtained using a biopsy puncher and these gels were placed into wells of a 48-well microplate. 300 μL of PBS was added into wells, incubated at RT, and at timepoints: 0, 4, 12, 24, 74, and 168 hours (or at 0, 0.2, 0.5, 1, 3.1, and 7 days), liquid samples were collected and replaced by fresh PBS. Spectral scans of these specimens were conducted using the NanoDrop spectrophotometer (Thermo Fisher Scientific, Waltham, MA) at 190 to 200 nm with PBS as a blank. Chemicals that can potentially leach out of the PEG-GLY gels (PEGDA, GLY, TEMED, and APS) were tested as well. The absorbance values were computed as the difference between samples and the blank PBS at 193 nm. GLY concentration was determined using the linear trendline of the glycerol standard curve. Mass release fraction was computed by dividing the released glycerol mass with the initial mass. A scatterplot of cumulative mass release versus time was then generated and a saturation curve fitted using least-squares regression in MATLAB with the equation:

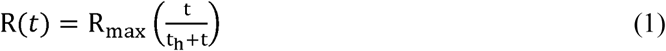

where R = cumulative mass release fraction (function of t), t = time, R_max_ = 100% (maximum release), t_h_ = saturation equation constant = time to release half the load.

### F. Bioactive glass synthesis and incorporation into gels

Bioactive glasses (BGHU1 and control Bioglass 45S5 (BG45S5)) were fabricated based on the modified protocol of Greenspan and Hench.^25^ Briefly, 50 g total of chemical powders: SiO_2_, Na_2_O, CaO, and P_2_O_5_ were homogenously mixed into a ceramic crucible (Engineered Ceramics, Gilberts, IL) and melted in an electric kiln at 1288 °C for 4 hours. The theoretical calcium to phosphorus mole ratios (n_Ca_/n_P_) of BGHU1 and BG45S5 were 4 and 5, respectively. Using metal tongs, the liquified glass was poured into new crucibles in another kiln set at 460 °C, incubated for 8 hours (annealing), and allowed to cool to room temperature for 24 hours. The synthesized glasses were removed from the crucible, and ground into fine particles with a mortar and pestle. PEG (20% PEGDA)-GLY (30%)-bioglass gels were made with 5% (m/V) bioactive glass particles using chemical crosslinkers. After the gelation process, additional 5% bioglass was added on the gel surface.

### G. Blood coagulation test

To investigate the potential host response effect of bioactive glass additives on gels, a clotting assay was performed. PEG-GLY-bioglass gels (PEG-GLY-BGHU1 and PEG-GLY-BG45S5) were made at 1.5 mL each into wells of a 12-well microplate. Control groups: no gels (surface of the poly(styrene) plate), PEG only, and PEG-GLY gels were also included. Samples were quickly rinsed with PBS to remove excess and unreacted reagents. Cat bloods in EDTA tubes collected from animal shelters with an approved Animal Care and Use Committee (ACUC) protocol from Dr. John Foy (Bobbi and the Strays, Freeport, NY) at 100 μL were mixed with 1 M CaCl_2_, then immediately added onto the test gel substrates. Blood coagulation time was recorded and normalized over the surface area of blood contact.

### H. Addition of alginate and laponite in hydrogels

To potentially improve the strength properties of PEG-GLY-BGHU1 constructs, alginate (ALG) and laponite (LAP) biomaterials were added. Specifically, PEG-GLY-BGHU1-ALG- LAP gels were made by mixing 20% PEGDA, 30% GLY, 5% BGHU1, 2.5% (m/V) ALG and 2.5% (m/V) LAP, then chemically crosslinked by 1% TEMED and 1 mg/mL APS. After curing for 15 minutes, gels were soaked in 1 M CaCl_2_ solution overnight to polymerize the alginate components. Compression until failure test was performed and properties obtained.

### I. Cyclic compression loading

Bigger PEG-GLY-BGHU1-ALG-LAP cylindrical gels were fabricated (at d = 50 mm and h = 10 mm) to model a human intervertebral (IV) disc between vertebrae lumbar 3 and 4 (L3 and L4). L3 and L4 vertebral bodies were simplified, modeled as cylinders that slightly taper in the middle, and three-dimensional (3D)-printed using a green poly(lactic acid) (PLA) filament (MakerBot, Brooklyn, NY). The hydrogel “IV disc” was positioned between PLA “vertebrae” and placed on the platen of Instron 3345. The Bluehill 3 software was configured for cyclic loading at maximum compressive strain of 4% (at 0.1 mm/s) for 100 cycles of compression and relaxation. Stress-strain curves were plotted. Dissipated energy density was calculated as the area of the hysteresis loop using MATLAB. A power equation was fitted using the formula:

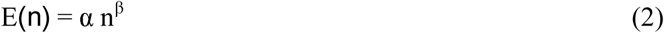

where E = dissipated energy density (function of n), n = cycle number, and α and β = constants.

### J. Statistical analyses and softwares

Experimental samples were performed in triplicates (n = 3). Values were reported as mean ± 1 standard deviation. Student’s t-test and one-way analysis of variance (ANOVA) with multiple comparison (Tukey’s) tests were performed in MATLAB at 5% probability (p) of type I error. Chemical structures of plasticizers were drawn using ChemSketch (Advanced Chemistry Development, Toronto, ON, Canada).

## III. RESULTS AND DISCUSSION

### A. Compression properties of bulk PEG gels with GLY, PEG, PRO, and SOR

Relative displacements and compressive loads (forces) were converted to true compressive strain and stress values, respectively, and plotted to obtain the elastic modulus at compression or compressive modulus (E_c_), yield, and ultimate strain and stress properties [Fig. 1].

**Fig. 1.**
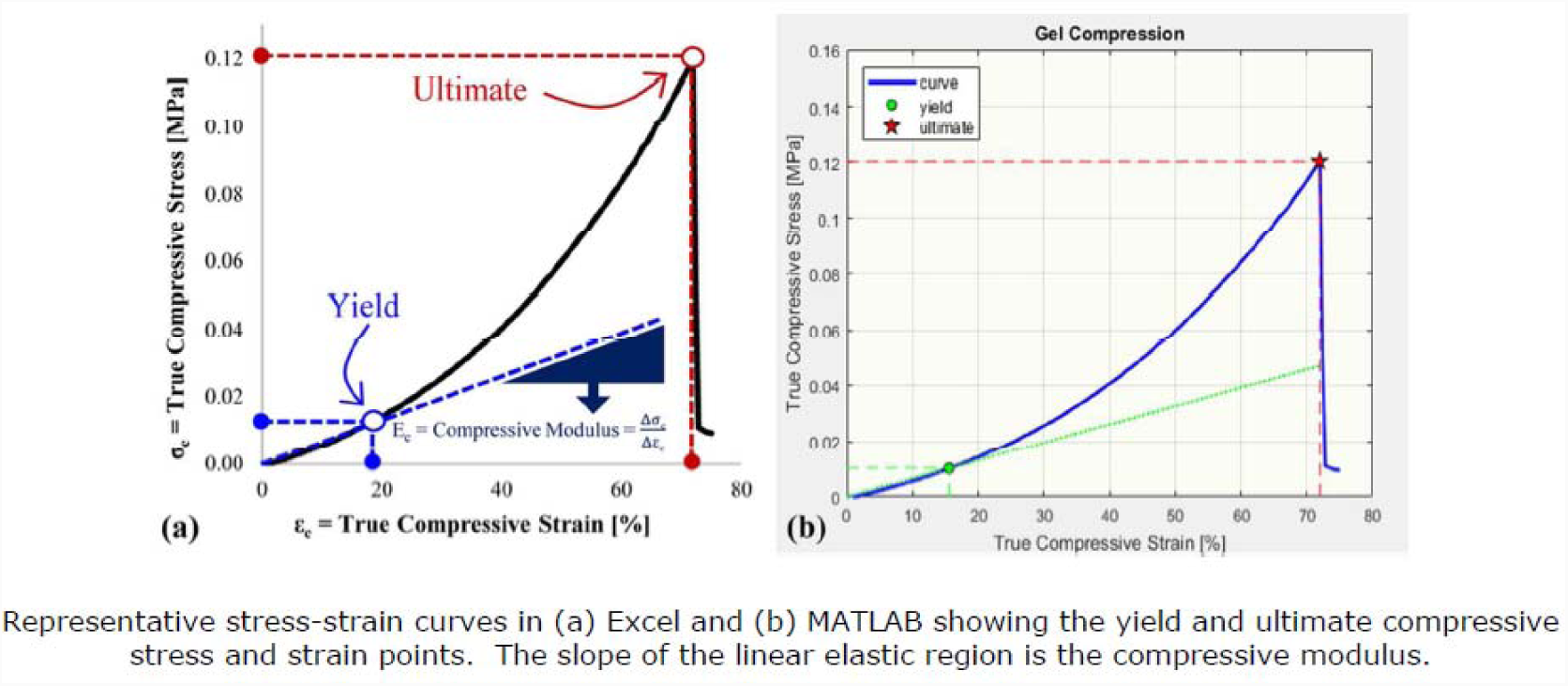

The shapes of the stress-strain curves of PEG gels (using 20% PEGDA) at varying plasticizer saturation concentrations changed expectedly; steepness of slopes increased then decreased [Fig. 2(a) showing PEG-GLY gels]. Most PEG gels made via photochemical polymerization did not fracture during the compression test. Hence, only the E_c_, yield stress (σ_c-y_), and yield strain (ε_c-y_) data were extracted from the stress-strain scatterplots. Plots of these properties versus plasticizer saturation [Fig. 2(b)-2(d)] revealed some patterns. The E_c_ values (slopes of the linear elasticity region) generally increased then decreased from 0 to 100% saturation [Fig. 2(b)] corresponding to plasticizing and anti-plasticizing effects^26^, respectively. PEG gels with 60% saturated SOR (or 750 mg/mL × 60% = 450 mg/mL SOR (Table I)) produced the highest E_c_ at 1.5 MPa. The staggered conformations of sorbitol^27^ possibly got entangled within the PEG matrix (at 60% saturation) leading to the highest stability and elasticity among the tested plasticizers. In other studies, sorbitol plasticizer also increased the strength properties of structural proteins films.^28, 29^ Glycerol, propylene glycol, and PEG 8000 (PEG with MW = 8000) also exhibited plasticizing effects at low saturation levels: 25%, 14%, and 50%, respectively [Fig. 2(b)].

**Fig. 2.**
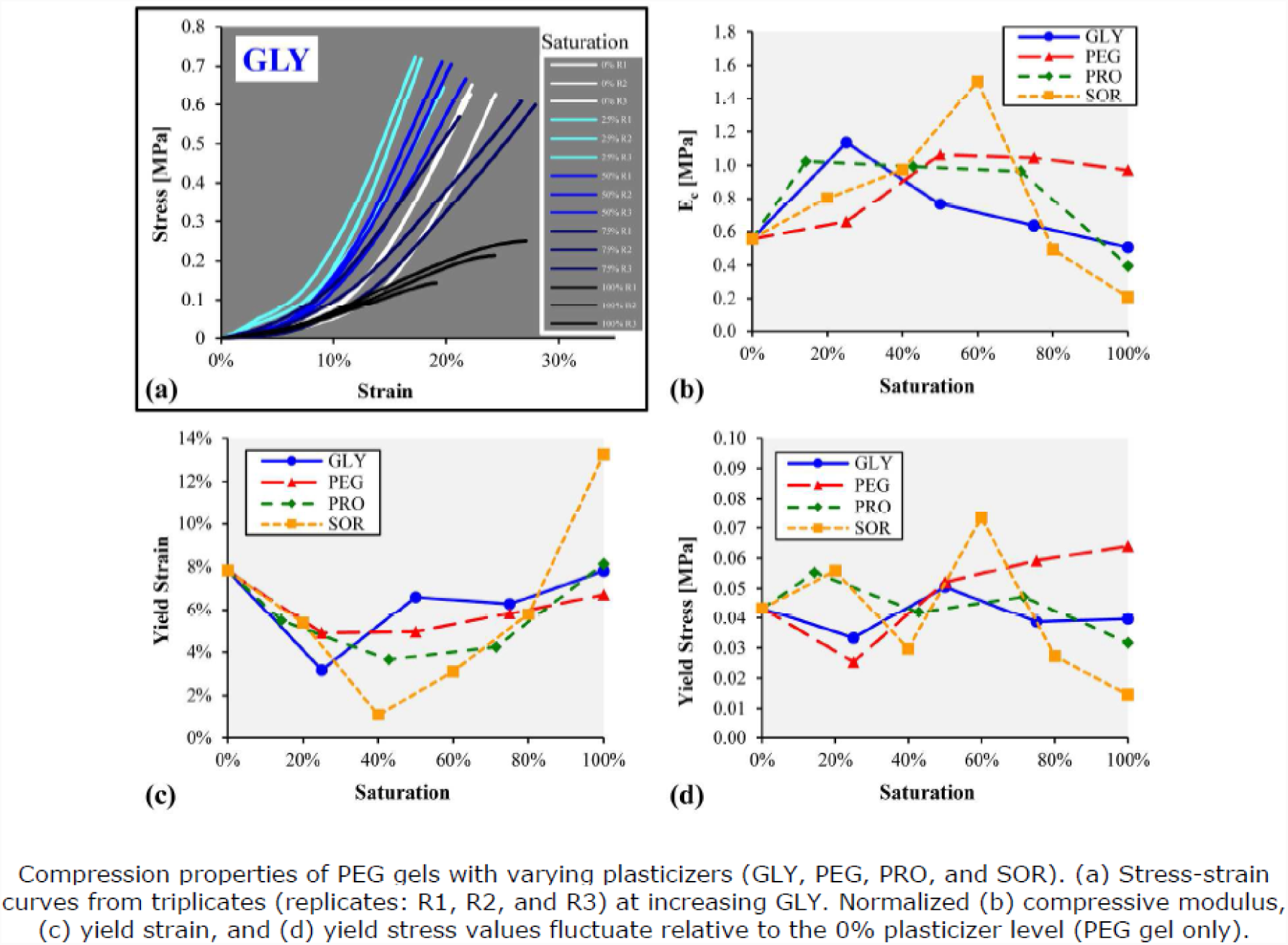

Addition of small amounts of plasticizers decreased the gel flexibility (decrease in ε_c-y_) but reverted to original values (of 8% strain) at plasticizer full saturation levels except for the gel with 100% saturated SOR that increased the ε_c-y_ to about 13% [Fig. 2(c)]. Compressive yield stresses erratically fluctuated at different plasticizer quantities [Fig. 2(d)]. Gels with 450 mg/mL SOR recorded 0.074 MPa = 74 kPa of σ_c-y_. Interestingly, the PEG-based gels with 75% saturated GLY (or 0.4 mL/mL × 75% = 0.3 mL/mL = 30% GLY) had E_c_, ε_c-y_, and σ_c-y_ values similar to the gels with no plasticizers, perhaps due to the balance between plasticizing and anti-plasticizing behavior of glycerol at the 30% concentration.

### B. Effect of temperature and pressure on gel properties

Chemical polymerization was utilized over the photochemical method for better control of gelation, especially for relatively thick gels. The chemically-crosslinked PEG-GLY (20% PEGDA with 30% GLY) gels fabricated in a heated die [Fig. 3(a)] showed an increase in opacity as temperature increased [Fig. 3(c)], consistent with the response of poly(N-isopropylacrylamide) (PNIPA) in which lower fabrication temperatures produced translucent gels while those at higher temperatures turned opaque.^30^ Temperatures > 80 °C resulted in damaged gels. The measured compression properties generated noticeable trends. Batch variability was small [Fig. 3(d)-3(e)], suggesting uniformity and repeatability of the chemical gelation process. ANOVA tests revealed significant differences among temperature groups for all the mechanical properties except for the ultimate compressive stress (p = 0.113). Gels fractured at 199 ± 48 kPa of compressive pressure. PEG-GLY gels made at 37 °C had significantly higher yield stress (at 52 ± 11 kPa) compared to the other temperature groups (p ≤ 0.0308) [Fig. 3(d)]. It was observed that as temperature increased, the compressive strain values also increased [Fig. 3(e)]. Gels polymerized at 80 °C were the most flexible, with yield and ultimate strains of 19 ± 2% and 70 ± 2%, respectively. Additionally, these gels (at 80 °C) produced significantly (p ≤ 0.0102) lower E_c_ at 89 ± 39 kPa compared to the other temperature groups at 345 ± 63 kPa. Heating increased the molecular motion and randomness of PEG chains’ orientation, producing increased light absorbance (opacity) and spring^31^ behavior.

**Fig. 3.**
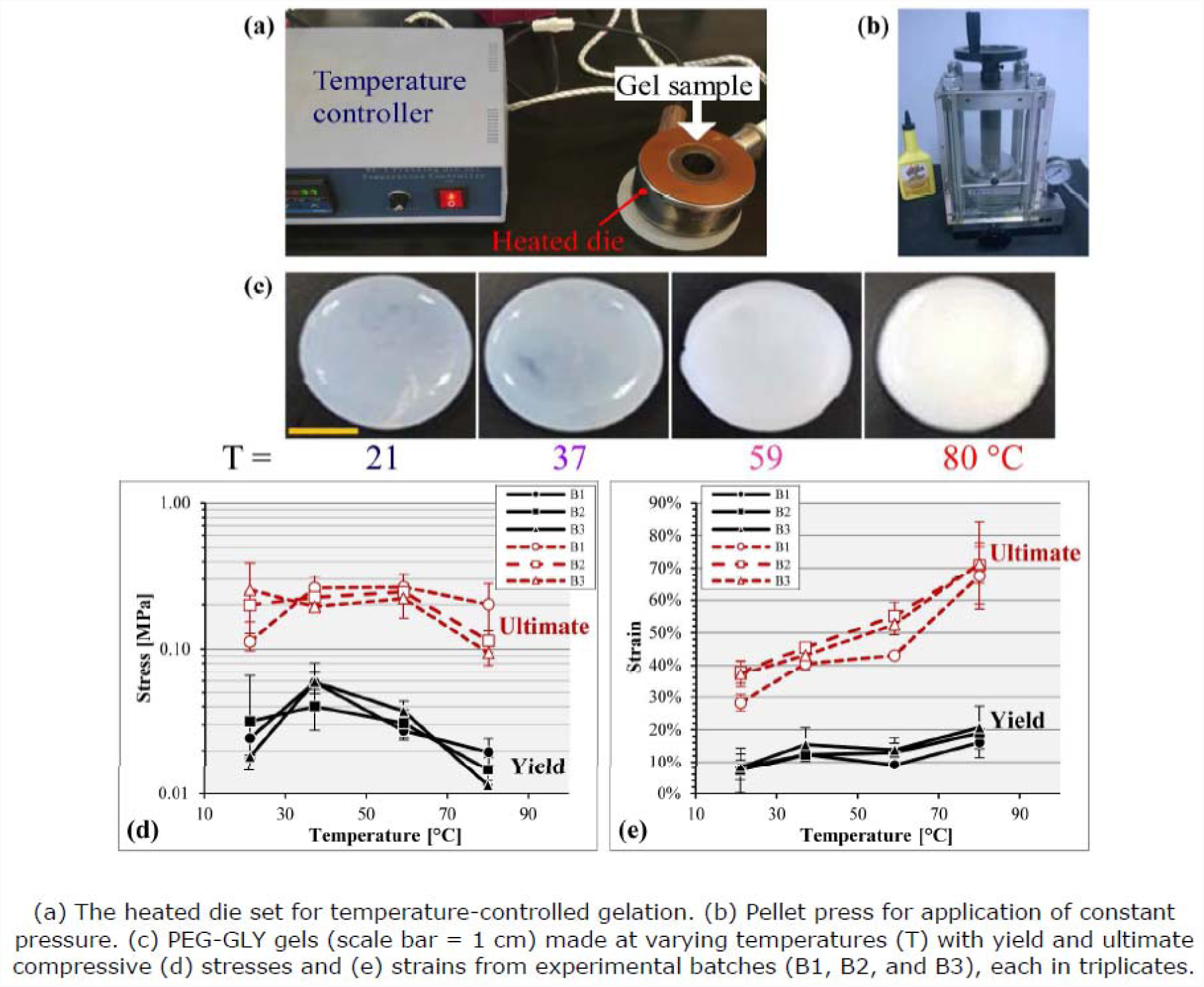

Application of constant gauge pressure of 90 MPa using a pellet press [Fig. 3(b)] at room temperature produced PEG-GLY gels that had statistically-lower E_c_ (p = 0.0112) at 36 ± 2 kPa and σ_c-y_ (p = 0.0228) at 5 ± 2 kPa compared to those made at ambient pressure. The incidence of untestable gels were high due to the die set leakiness at the pressurized condition; thus, the pellet press system needs improvement.

Pairwise comparison of PEG gels revealed that at 80 °C, those with GLY had statistically lower ultimate compressive strength (136 ± 57 kPa vs. 310 ± 42 kPa; p = 0.0194) and compressive modulus (89 ± 39 kPa vs. 294 ± 55 kPa; p = 0.0329) than without any plasticizer [Fig. 4(a)]. However, at 37 °C, the addition of GLY led to PEG gels with increased yield stress (p = 0.031) and yield strain (p = 0.0216) [Fig. 4(b)]. GLY also generated gels with higher (p = 0.0402) ultimate compressive strain at 21 °C [Fig. 4(b)].

**Fig. 4.**
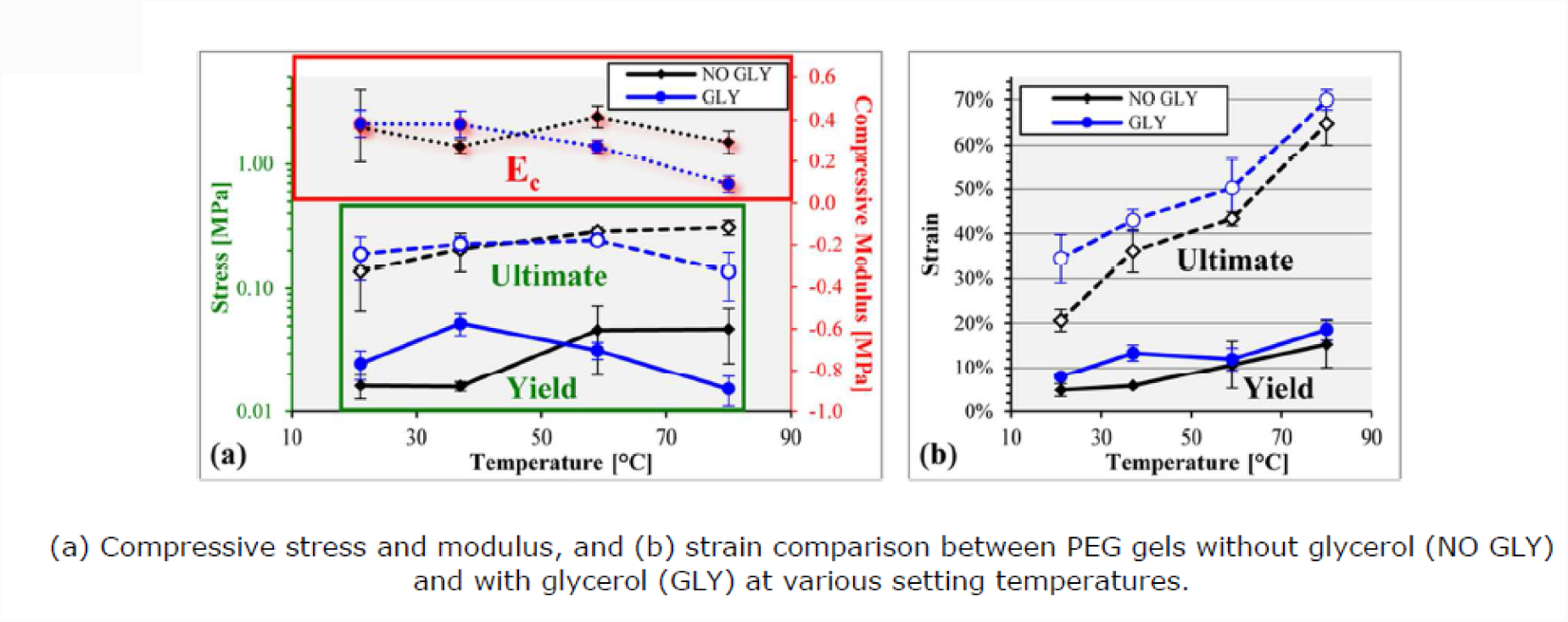

### C. Chemical versus photochemical gelation

PEG hydrogels were crosslinked and polymerized using chemical (APD and TEMED) and photochemical reactions (I2959 and light at 254 nm). It was determined through mechanical testing that the compressive modulus of PEG gels with GLY were significantly greater (p = 0.0387) when photopolymerized than chemically gelled [Fig. 5]. Moreover, photochemical PEG-only gels had statistically higher yield strain (p = 0.0143) and stress (p = 0.0296) compared to those made using chemical crosslinking [Fig. 5].

**Fig. 5.**
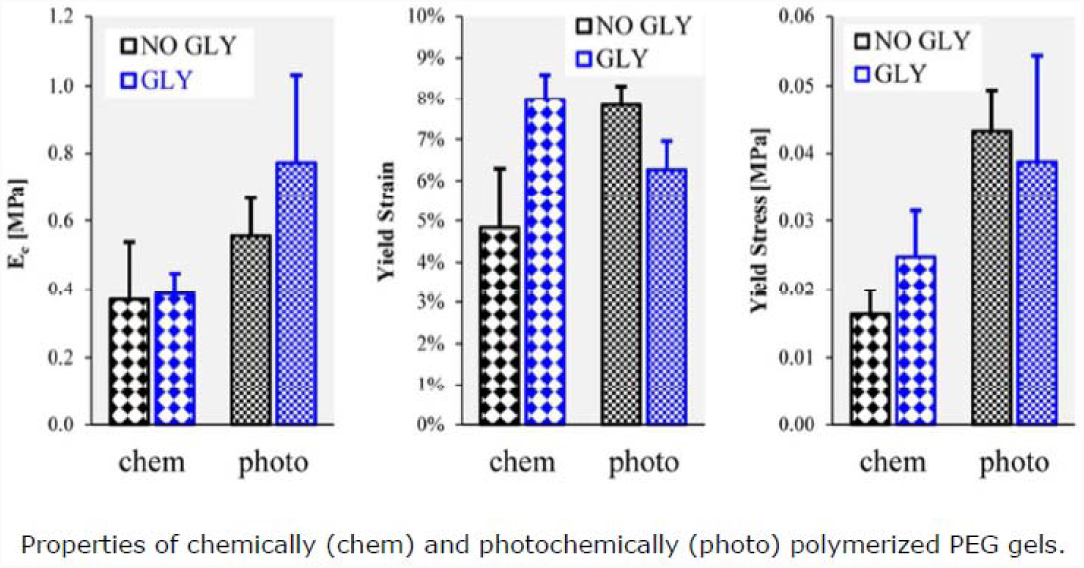

### D. Stability of glycerol in PEG hydrogels

The subsequent experiments employed PEG gels with glycerol as the chosen plasticizer. Accordingly, glycerol’s retention within the PEG matrix was evaluated. The glycerol standard curve followed a linear trendline [Fig. 6(a); r^2^ = 0.96629] with the equation: C = (A – 0.8137)/1.6809, where C = GLY concentration and A = absorbance at 193 nm. At 193 nm, GLY generated the highest positive signal difference compared to water, PBS, PEGDA, TEMED, and APS (possible substances in the collected liquid). Based on this linear equation, the mass release fraction values were calculated from light absorbances. It was found that the cumulative release of glycerol out of the PEG-GLY gels [Fig. 6(b)] was time-dependent and behaved according to the saturation equation (r^2^ = 0.94571): R = t/(t_h_ + t), where t_h_ was determined to be 0.9535 day = time needed to release half the amount of GLY from the gel. At the 1-week experimental endpoint, PEG-GLY gels had diffused out 89 ± 39% of the original glycerol mass load, demonstrating that glycerol was not covalently-bound to the PEG matrix and leached out quickly into the PBS medium. It is thus advised that PEG-GLY gels are used in minimal liquid environment to prevent glycerol release.

**Fig. 6.**
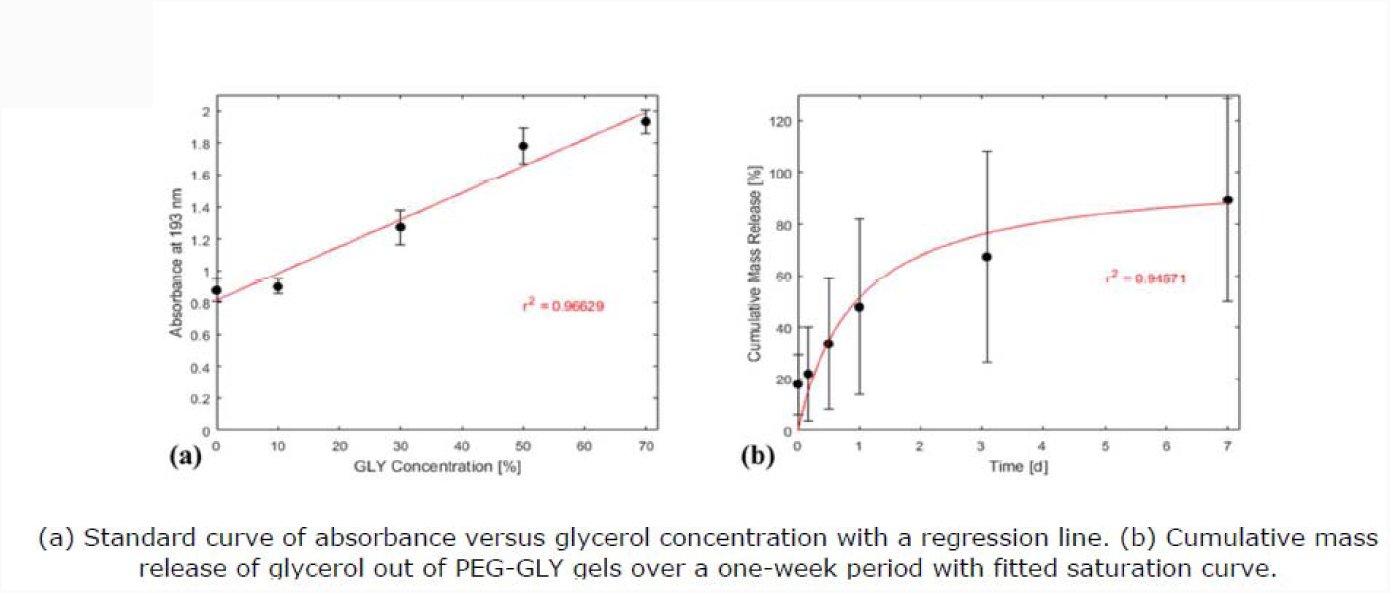

### E. Blood clotting on gels with bioactive glass particles

Using kilns [Fig. 7(a)] for melting and annealing, bioglass particles were successfully synthesized [Fig. 7(b)], then incorporated into PEG-based gels. Blood coagulation on gel surfaces [Fig. 7(c)] revealed that PEG-GLY-BGHU1 gels (PEG-GLY hydrogels with BGHU1 bioactive glass particles on exposed surfaces) induced significantly faster clotting time per unit area compared to poly(styrene) surfaces (no gel; p = 0.0207) and to PEG only (p = 0.0334) gels [Fig. 7(d)]. The blood coagulation on PEG-GLY-BGHU1 was also lower at 63 ± 11 s/cm^2^ versus on PEG-GLY at 81 ± 21 s/cm^2^, but statistically not significant (p = 0.7721). Gels with bioglass (PEG-GLY-BG45S5 and PEG-GLY-BGHU1) activated similar rates of blood clotting (p = 0.9998), implying that the new bioactive glass mixture (BGHU1) and the control Bioglass 45S5 were alike in their hemostatic capabilities. Other bioactive glass formulations had also been shown to increase hemostasis and *in vitro* blood coagulation.^32, 33^ Faster blood clotting time may suggest better host tissue integration but ultimately requires actual *in vivo* implantation experiments.

**Fig. 7.**
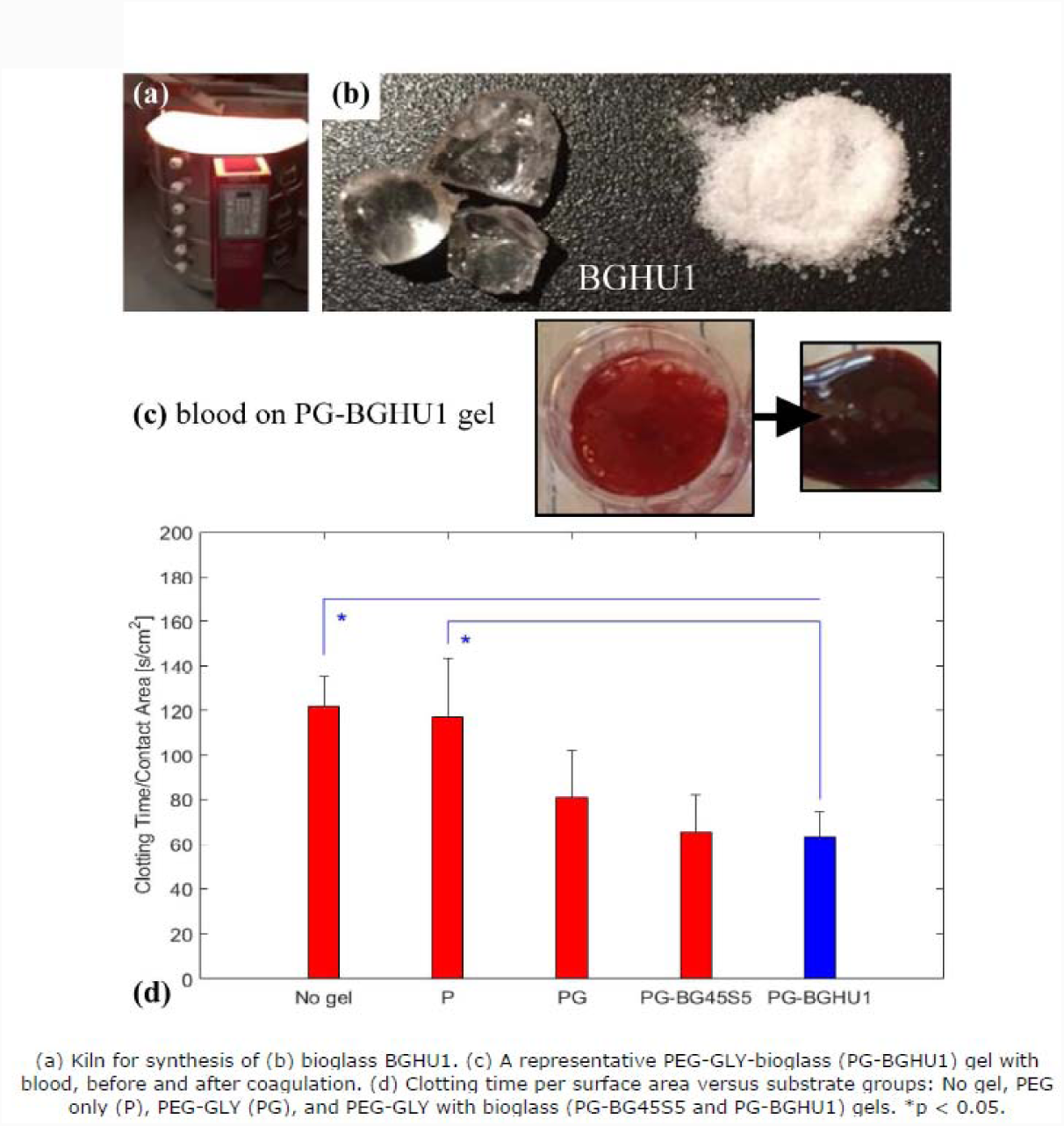

### F. Biomechanical properties of gels with bioglass, alginate, and laponite additives

PEG-GLY-BGHU1 gels had compressive modulus, yield strain, and yield strength results statistically-similar (p > 0.05) to gels (PEG-GLY) without the bioglass particles (Table II). Bioactive glass particles were only added on the gel surfaces, hence, did not alter the bulk properties. PEG-GLY hydrogels with added alginate (at 2.5%) and laponite (at 2.5%) constituents (PEG-GLY- BGHU1-ALG-LAP) produced higher E_c_ (p ≤ 0.0431), ε_c-y_ (p ≤ 0.014), and σ_c-y_ (p ≤ 0.0187) measurements compared to the groups without ALG and LAP (Table II). The ultimate strain and stress points of PEG-GLY-BGHU1 gels were determined to be 34 ± 1% and 254 ± 31 kPa, respectively. In another study using PEG gels, the presence of alginate and laponite additives also increased the mechanical strength of gels, particularly the fracture toughness or ultimate compressive strength due to the supplemental alginate network formation and reinforcing laponite load-bearing structures.^21^ The 79-kPa yield strength of the PEG-based gel IV disc model was below the 100 kPa computed biomechanical tolerance of a native lumbar IV disc of an average man in supine position^34^, therefore the gel assembly needs further research and development.

**Table II.**
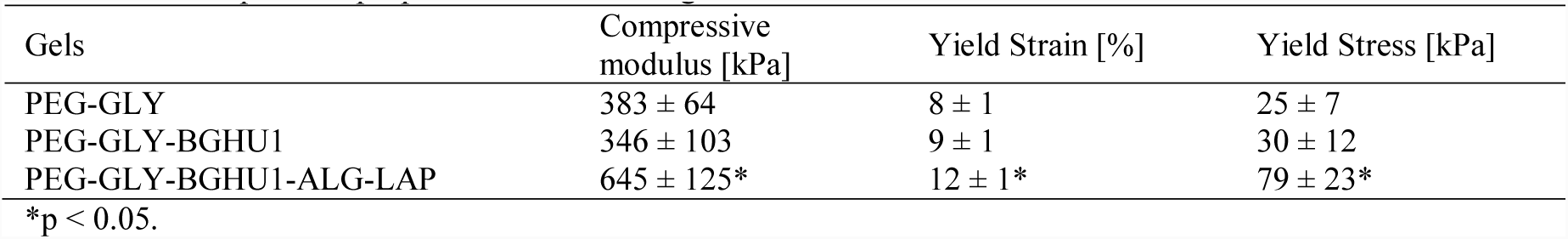
Compressive properties of PEG-GLY gels.

| Gels | Compressive modulus [kPa] | Yield Strain [%] | Yield Stress [kPa] |
| --- | --- | --- | --- |
| PEG-GLY | 383 ± 64 | 8 ± 1 | 25 ± 7 |
| PEG-GLY-BGHU1 | 346 ± 103 | 9 ± 1 | 30 ± 12 |
| PEG-GLY-BGHU1-ALG-LAP | 645 ± 125* | 12 ± 1* | 79 ± 23* |

\*p < 0.05.

PEG-GLY-BGHU1-ALG-LAP gels were geometrically modeled as cylindrical lumbar spine IV discs [Fig. 8(a)] and tested for cyclic compression [Fig. 8(b)] at an initial compressive strain of 4% (determined to be within the linear elasticity region of the gel). Hysteresis and creep responses were observed during cycles of compression (loading) and relaxation (unloading) [Fig. 8(c)], with similarities to behavior of actual IV discs during cyclic compression.^35, 36^ Stress-strain curves were found to decrease in magnitude at increasing number of cycles (n) [Fig. 8(d)]. Consequently, hysteresis behavior and dissipated energy density (E = area within the hysteresis loop) also decreased following a power regression (r^2^ = 0.97) with the equation: E = α nβ, where α = 4.5102 J/m^3^ = dissipated energy density at the first cycle and β = –0.4069 = exponent [Fig. 8(e)], indicative of elastic recovery^37^ exhibited by biological tissue specimens.^38, 39^

**Fig. 8.**
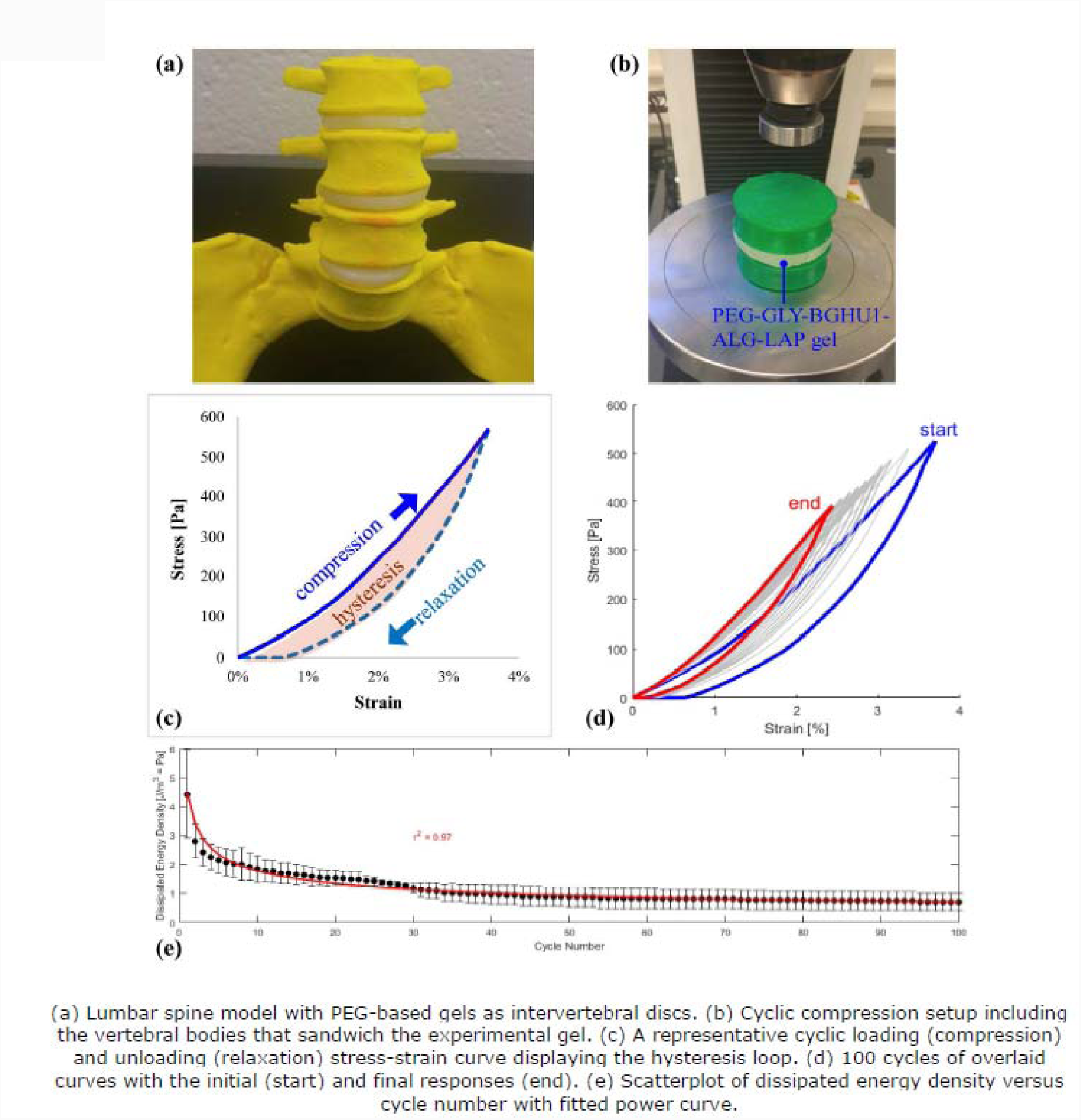

## IV. CONCLUSIONS

This research study showed improvements of compressive strengths of 20% PEG gels through the use of GRAS plasticizers (sorbitol, glycerol, propylene glycol and PEG 8000) at relatively low saturation concentrations. Photochemical reaction proved to be the superior method of gelation over chemical for creation of stronger gels; but chemical means provided better control and product uniformity. The gelation temperature of 37 °C made constructs (PEG with 30% glycerol) with the highest yield strength among the other temperature groups. Gels synthesized at ambient pressure, with the addition of alginate and laponite, led to greater increase in the mechanical performance. These gels with surface bioglass particles, modeled to cylindrical lumbar intervertebral discs, demonstrated decent biomechanical responses. Additional experimentation and development activities are needed for translational applications.

## ACKNOWLEDGMENTS

The authors would like to thank the members of the Bioengineering Materials Lab including Daniel Foyt and Hazel Consunji for technical assistance, manuscript proofreading, and discussion, Tyler Lavertu for gel fabrication under extreme pressure, Paul Chaleff and Bethany Dill for help with bioglass synthesis, John Foy for providing the animal blood samples, Meir Wieder for the Excel Visual Basic for Applications (VBA) coding, Pierre Llanos and Robert Cerro for 3D printing assistance, and Jacqueline Scarola and Lori Castoria for help with reagents purchasing. Support for this study was provided by Hofstra University internal research funds.

